# Metacommunity stability and persistence for predation turnoff in selective patches

**DOI:** 10.1101/2021.10.26.465836

**Authors:** Dweepabiswa Bagchi, Ramesh Arumugam, V K Chandrasekar, D V Senthilkumar

## Abstract

Predation as an important trophic interaction of ecological communities controls the large-scale patterns of species distribution, population abundance and community structure. Numerous studies address that predation can mediate diversity and regulate the ecological community and food web stability through changes in the behaviour, morphology, development, and abundance of prey. Since predation has large effects on persistence and diversity, the local loss or removal of predation in a community can trigger a cascade of extinctions. In ecological theory, the effect of predation removal has been well studied in foodwebs, but it remains unclear in the case of a spatially distributed community connected by dispersal. In this study, the interaction between local and spatial processes is taken into account, we present how a predation turnoff in selective patches affects the stability and persistence of a metacommunity. Using a simple predator-prey metacommunity with a diffusive dispersal, we show the impact of predation on synchronized, asynchronized and source-sink dynamics. Our results reveal that predation turnoff in very few patches alters a perfectly synchronized oscillatory state into multicluster states consisting of various patterns. In a source-sink behaviour, predation turnoff in a source patch reduces the number of sink patches and changes the clusters. In general, predation turnoff in a finite number of patches increases the number of clusters through asynchronized (inhomogeneous) states, whereas predation turnoff in a larger number of patches can lead to the complete extinction of predators. Typically, there exists a critical number of patches below which the predation turnoff results in asynchronized states and above that predation turnoff leads to a synchronized state in prey population with complete extinction of predators. Further, our results identify the network configurations that exhibit a unique number of clusters. Moreover, prey density from the patches where predation is absent goes to a saturating state near the carrying capacity. Thus, this study stresses that predation turnoff in selective patches acts as a stabilizing mechanism that can promote metacommunity persistence.

## 1. Introduction

Various interactions such as competition, predation, mutualism, dispersal, etc., shape the structure and functioning of ecosystems (Brewer, 1979; Menge and Olson, 1990; Loreau et al., 2002). Predators are ubiquitous in natural environments, and their effects on local communities are well recognized. Numerous studies address that predation at different trophic levels of ecological communities or foodwebs controls the large-scale patterns of species distribution, population abundance and community structure (Menge and Sutherland, 1987; McCann et al., 1998; Chalcraft and Resetarits Jr, 2003). Also, predation can mediate the coexistence of many species in the ecological community despite that it can reduce the prey population densities (Paine, 1966; Leibold, 1996; Ryberg et al., 2012). Through the ‘top-down effect’, predation can regulate the ecological community and foodweb stability (Carpenter and Kitchell, 1996; Leibold, 1996; Pace et al., 1999; Duffy, 2002). Subsequently, in spatially distributed communities, predator and its dispersal alter the diversity at local and spatial scales (Shurin and Allen, 2001; Chase et al., 2009; Ryberg et al., 2012; Johnston et al., 2016). Moreover, predators can cause changes in the behavior, morphology, development, and abundance of prey (Werner and Peacor, 2003; Brown and Kotler, 2004). Since predation has large effects on persistence and diversity (Paine, 1966), the local loss or removal of a predator in a community can trigger a cascade of secondary extinctions (Quince et al., 2005; Borrvall and Ebenman, 2006). The effect of predation removal has been well studied in foodwebs, but it remains unclear in the case of a spatially distributed community connected by dispersal. Indeed, predator removal in spatially distributed patches can have far-reaching consequences for the structure and functioning of ecosystems. Hence, in this study, we address how predation turnoff in selective patches affects the metacommunity stability and persistence.

Empirically, drastic and detrimental changes to metacommunity following loss of predators have been proved in numerous cases (Crooks and Soulé, 2005; Pech et al., 1992; Ritchie and Johnson, 2009; Wallach et al., 2010). Indeed, predators have an adverse effect on domesticated animals (Gusset et al., 2009; Mishra, 1997; Oli et al., 1994), prey species (Dalla Rosa and Secchi, 2007; Henschel et al., 2011; Weise and Harvey, 2005), and human populace (Dickman, 2010; Gore et al., 2005; Löe and Röskaft, 2004; Penteriani et al., 2016). This rather limited view of predators has led to the consideration of removal of predators as a successful management tool by different state, regional and private agencies (Reynolds and Tapper, 1996; Bergstrom et al., 2014). Poison baiting, trapping, hunting or culling have been different ways of implementation of these methods. Such methods have scientifically been shown to have much lesser efficacy than expected, sometimes leading to utter failure in community sustenance (Lennox et al., 2018). They are also some of the costliest endeavors undertaken by governments (Berger, 2006). The question of sustaining predator population without culling of predators is therefore to be considered seriously. From a dynamical perspective, it is similar to turning off the predation without changing the other time-based dynamics of prey and predators. In a single community approach, husbandry practices (Jackson and Wangchuk, 2004; Johnson and Wallach, 2016), fencing (Hayward and Kerley, 2009) and deterrent devices (e.g., fladry) (Ogada et al., 2003) have been used to address such a scenario. However, the question of efficacy of such methods for a metacommunity remains completely unexplored. This work addresses that exact issue, i.e., the effect of predation turnoff on the dynamics of a large prey-predator metacommunity as a whole.

As far as metacommunity dynamics are concerned, population synchrony is an important dynamics referring the coherent behaviour of population densities (Ranta et al., 1995; Bjørnstad et al., 1999; Lande et al., 1999). Synchrony at temporal and spatial scales has been studied in many ecological systems since it might affect the stability of system by elevating a high risk of extinction (Heino et al., 1997; Earn et al., 2000). Typically, three major mechanisms induce the synchronized behavior: climatic disturbances (i.e. the Moran effect), trophic interactions (i.e., predation), and dispersal among populations (Bjørnstad et al., 1999; Koenig, 1999; Ims and Andreassen, 2000; Liebhold et al., 2004). In particular, the Moran effect addresses that correlation in environmental conditions can cause synchronous fluctuations over time (Moran, 1953; Ranta et al., 1997; Ripa, 2000; Engen and Sæther, 2005). Predation as a trophic interaction with populations of other species can inhibit a synchronizing effect with either prey populations at a lower trophic level or with species at a higher trophic level (Ims and Steen, 1990; Korpimäki and Norrdahl, 1998; McCann et al., 1998; Satake et al., 2004; Liebhold et al., 2004). Among three mechanisms, dispersal is a major one studied widely in spatially distributed communities. Dispersal among populations can destabilize communities by inducing correlated population dynamics across space resulting in spatial synchrony (Liebhold et al., 2004). On the other end, it can also reduce local extinction risk through asynchronized dynamics and stabilize the metacommunity thorough recolonization or rescue effects (Hanski, 1998, 1999; Jansen, 2001; Abbott, 2011). It is well known that dispersal is one of the stabilizing processes of spatially connected ecological systems (Briggs and Hoopes, 2004). Nevertheless, the combinations of dispersal, predation and environmental correlation have been used in numerous studies to understand the metacommunity stability (Lande et al., 1999; Kendall et al., 2000; Gouhier et al., 2010; Goldwyn and Hastings, 2011; Arumugam and Dutta, 2018).

The interaction between predation and dispersal (as local and spatial processes) has disproportionately large effects on the persistence (Holyoak et al., 2005; Howeth and Leibold, 2013; Johnston et al., 2016). Though dispersal can synchronize the populations, predator dispersal can lead the metacommunity into asynchronized dynamics through inhomogeneous states and source-sink dynamics (Amarasekare and Nisbet, 2001; Gravel et al., 2010). In general, metacommunity with asynchronized states and source-sink behavior has less risk of extinction and higher persistence as compare to their synchronized behavior. Predator is indeed a keystone species in spatially distributed communities. In such a scenario, predation control or removal even in some selective patches can have far-reaching consequences for the structure and functioning of ecosystems. Hence, in this study, we present how local and spatial processes (i.e., predation and dispersal) interact to produce patterns that reduce extinction risk of the metacommunity. In particular, we address (i) how predation turnoff in some selective patches affect the synchronized, asynchronized and source-sink dynamics of a metacommunity? (ii) how predation turnoff in the source patch affects the source-sink dynamics? and (iii) how patch-wise predation turnoff and even increasing number of patches in predation turnoff affect the metacommunity stability?

In the theory of metacommunities, coupled ecological oscillators with different network topologies, direct and indirect coupling schemes constitute an efficient framework to understand the stabilizing effects of dispersal (Goldwyn and Hastings, 2008; Holland and Hastings, 2008; Arumugam et al., 2015, 2019). To address the effect of predation turnoff, we use a simple predator-prey metacommunity in a network of patches connected by a diffusive dispersal. By applying the predation turnoff in some patches, we study the metacommunity dynamics. Both synchronized, asynchronized and source-sink dynamics are taken into account in our analysis, we present how predation turnoff affects the stability and persistence of the metacommunity. Using the cluster size as a dynamical property of the metacommunity network, we present how patch-wise predation turnoff as well as predation turnoff in the increasing number of patches affect the stability of the system. Our results reveal that predation turnoff can alter the synchronized dynamics into asynchronized dynamics by manifestation of multi-cluster from a single cluster. In source-sink dynamics, the predation turnoff in the source patch reduces the number of sink and subsequently increases the cluster size. Further, our findings reveal the unique network configuration for predation turnoff to exhibit a specific cluster. Moreover, the prey density from the patch where the predation is turned off goes to a saturating population density near the carrying capacity. Thus, predation turnoff in selective patches increases asynchrony in metacommunity, which in turn promotes persistence.

In the next section, a schematic representation of predation turnoff and a predator-prey metacommunity model are described in detail. In the ‘Results’ section, we present the metacommunity dynamics for predation turnoff using synchronized and asynchronized dynamical states. Further, we present how patch-wise predation turnoff affect the metacommunity stability. Subsequently, we present how the predation variation affects the dynamics. Finally, in the ‘Discussion’ section, we address the ecological importance of our results and their consequences. In the supplementary material, we include the dynamics of predation turnoff considering other network topologies such as fully-connected network and a random network.

## 2. The Model

We use a predator-prey metacommunity in a network of *n* patches in which each patch is described by the Rosenzweig–MacArthur model (Rosenzweig and MacArthur, 1963). The local interaction between the prey (*V*) and the predator (*H*) within the patch is described by logistic growth and a type II functional responses, whereas the spatial interaction between the patches is described by a simple diffusive dispersal using a ring network topology (Jansen, 2001; Goldwyn and Hastings, 2008). Hence, in each patch (*i*), the dynamics of prey (*V*) and predator (*H*) are described by the following differential equations:

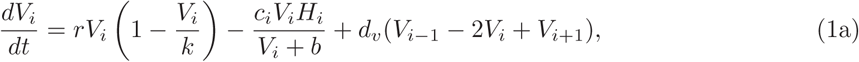

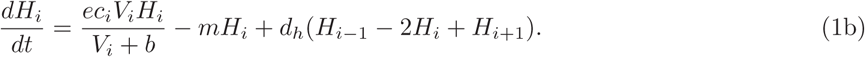

Here *i* = 1, 2, …, *n* and *n* denotes the total number of patches. The parameters of the prey are intrinsic growth rate (*r*), carrying capacity (*k*), and the half saturation constant (*b*), whereas the parameters of the predator are predation rate (*c*), predation conversion efficiency (*e*), and the mortality rate (*m*). In the coupling function, *d*_*v*_ and *d*_*h*_ denote the prey and predator dispersal strength, respectively. The ring-type coupling describes the connectivity between nearest neigboring patches.

The main aim of this study is to understand how predation turnoff affects the metacommunity stability. Using a schematic representation for a metacommunity network, we describe the predation turnoff in Fig. 1. Here circles denote the patches consisting of predator-prey interactions, whereas the connectivity between patches is denoted by lines. In Fig. 1, the right and left panels describe the metacommunity network structure with and without predation turnoff, respectively. In the left panel, the patches are identical when there is no predation turnoff. In the right panel, we turnoff the predation in some patches represented by an orange circle. For the predation turnoff in the *i*-th patch, we set *c*_*i*_ = 0 in Eq. (1). For the other patches (where there is no predation turnoff), *c*_*i*_ ≠ 0 and remains a positive constant. Note that we do not change the connectivity between patches in Fig. 1, but we turnoff only the predation in some selective patches. In other words, only local interaction within the patch is changed, but spatial interaction remains unaltered in the network.

**Figure 1:**
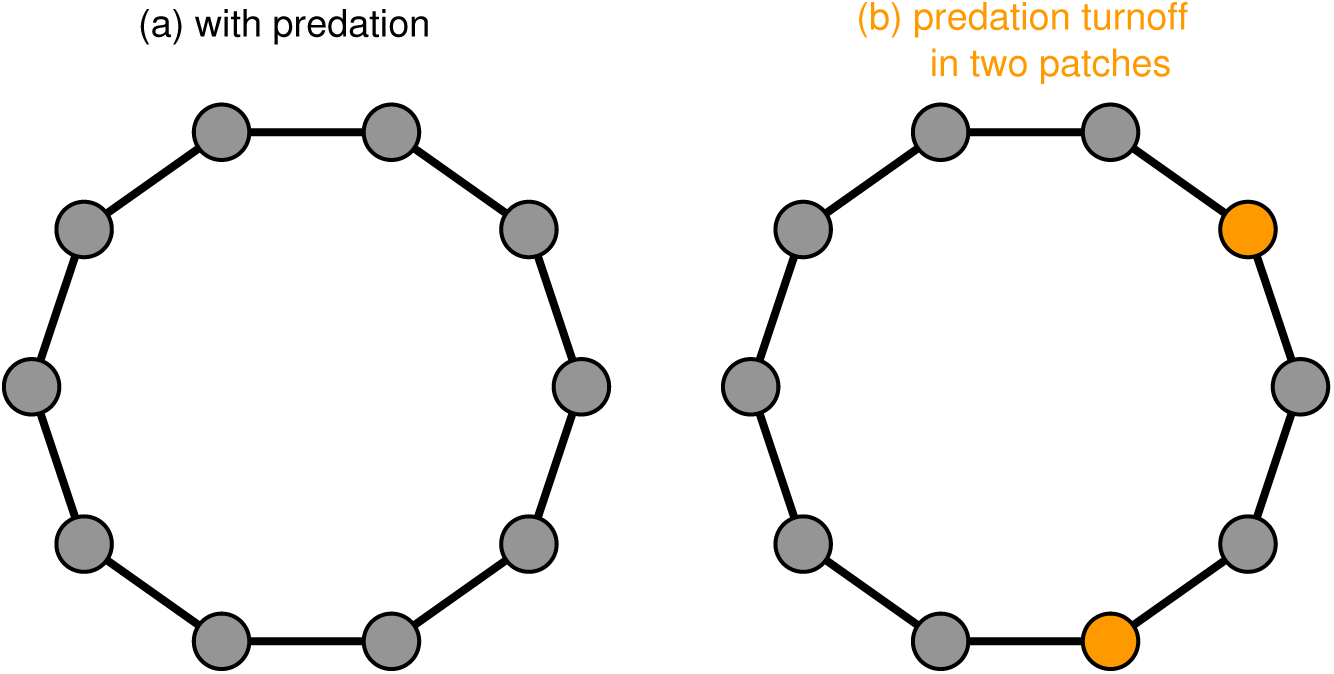
Schematic representation of predation turnoff in a metacommunity: (a) identical patches representing no predation turnoff in a ring network topology, (b) predation turnoff in two patches (highlighted in a color node). Here circles denote the patches consisting of prey-predator interaction, whereas lines denote the connectivity between the patches.

From the ecological perspective, the phenomenon of predation turnoff can be manifested as an antipredator adaptation mechanism developed through evolution in the prey species against predators (Lima and Dill, 1990; Caro, 2005). Few examples of such mechanisms are successful camouflage and schooling effect (Cott, 1940; Clark, 1974). Successful camouflage is when the prey seamlessly hides within its surrounding habitat without being seen. Schooling effect is the phenomenon where a large number of prey species elements commune together to form a large school, thereby diminishing the chance of one prey being singled out and predated.

## 3. Results

We begin our analysis by addressing the dynamics of the spatially distributed community when there is no predation turnoff. Numerous studies have dealt with prey-predator metacommunity dynamics using a simple diffusive dispersal (represented by a ring network topology in the left panel of Fig. 1) to understand the stability and persistence of spatially distributed ecological system (Jansen, 2001; Holland and Hastings, 2008). It is well known that dispersal can lead the metacommunity into synchronized and asynchronized dynamics. Specifically, metacommunity can exhibit homogeneous (synchrony) and inhomogeneous (asynchrony) spatially distributed population densities depending on dispersal rate. Often, metacommunity persistence can be addressed by the synchronized and asynchronized dynamics (Holland and Hastings, 2008). The former leads to a high risk of extinction, whereas the latter leads to more persistence (Briggs and Hoopes, 2004; Holland and Hastings, 2008).

Further, dispersal can prevent local extinction by exhibiting a source-sink dynamics (source and sink are characterized by high and low quality patch respectively). In metacommunity, source-sink behavior is often described by the inhomogeneous states among patches in which the population density is higher in some patches (source) and it is lower in other patches (sink). In this study, synchronized, asynchronized and source-sink dynamics are taken into account, we present how predation turnoff affects metacommunity stability and persistence using clusters. Cluster in this study refers to a group of patches showing a similar dynamical behaviour. For example, in 1-cluster case, all patches are in one group showing an identical behaviour. Similarly, in a 3-cluster case, three groups of patches are formed and patches from each group shows an identical population density. From numerous studies, ecological significance of clusters is well known, specifically on metacommunity persistence (Holland and Hastings, 2008). For example, population dynamis with 1-cluster has a high risk of extinction because it describes a perfectly synchronized behaviour, whereas multi-cluster (i.e., more than 1-cluster) has a less risk of extinction since it describes the asynchronized behaviour.

### 3.1. Asynchronized dynamics through predation turnoff

Using ten patches in the ring network topology, we analyse the predator-prey metacommunity dynamics. At first, we consider a simple case where the metacommunity exhibits a perfectly synchronized oscillation (i.e., 1-cluster) when there is no predation turnoff. Then, we turn off the predation in one or two selected patches (represented by a schematic in the left panel of Fig. 2 and the corresponding dynamics represented as time series in the right panel). Since we consider 1-cluster before predation turnoff, initial population density is patch-wise identical. The predation turnoff in only one patch alters the perfectly synchronized oscillation into a phase-synchronized oscillation with 6-cluster (see Fig. 2a).

**Figure 2:**
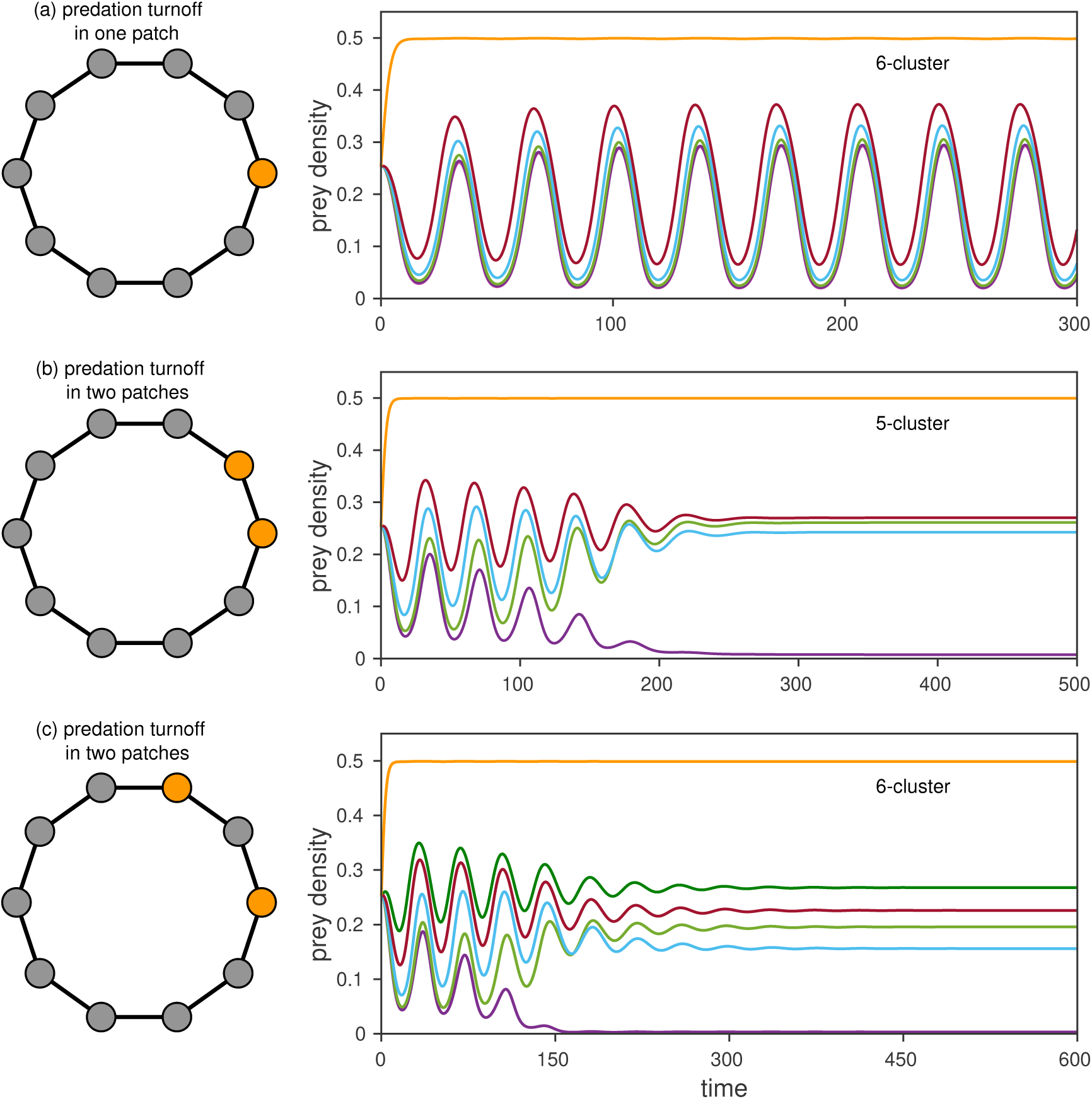
Metacommunity dynamics for predation turnoff in a finite number of patches: Considering a synchronized oscillatory state (1-cluster) in 10 patches, we show the schematic of predation turnoff in the left panel and the corresponding dynamics in the right panel. Metacommunity exhibits (a) 6-cluster for predation turnoff in one patch, (b) 5-cluster for predation turnoff in two consecutive patches, and (c) 6-cluster for predation turnoff in two patches (alternative to each other). Here the parameter values are *r* = 0.5, *k* = 0.5, *b* = 0.16, *e* = 0.5, *m* = 0.2, *d*_*v*_ = 0.001, and *d*_*h*_ = 1.3.

After removing enough transients, we compute the clusters using the time series of prey population density with error tolerance 10^−4^ (in this study, all the time series are computed using the fourth-order Runge-Kutta method). The prey density from the patch where the predation is turned off always reaches the highest density near the carrying capacity. Indeed, it forms a cluster showing oscillatory behavior albeit with extremely low amplitude. One can choose any random patch to remove the predation, but we observe a similar behaviour as in Fig. 2(a). Due to the symmetric behaviour of the ring topology, predation turnoff in one random patch always leads to 6-cluster with a phase synchronization. From an ecological perspective, this 6-cluster in the metacommunity has less risk of extinction as compared to the 1-cluster. Thus, predation turnoff in one patch reduces the extinction risk by increasing the number of clusters.

Now we analyse the dynamics for predation turnoff in two patches using the synchronized oscillatory state as our initial dynamics. Two patches can be chosen in 45 ways among the ten patches and we present two specific cases. At first, if two patches are chosen successively in the ring network (i.e. patch index *i* and *i* + 1), the metacommunity exhibits a stable steady state with a 5-cluster dynamics (see Fig. 2b). Indeed, the predation turnoff in two patches changes the synchronized 1-cluster oscillatory state into a 5-cluster steady state. In the second case, if two patches are chosen with one alternative to the other (i.e., the patch index *i* and *i* + 2), then the metacommunity exhibits a 6-cluster (see Fig. 2c). As mentioned earlier in one patch predation turnoff, the prey density from the patches where the predation is turned off goes to a saturating state with the highest density near the carrying capacity. Typically, two patch predation turnoff increases the number of clusters and also changes the dynamics from oscillatory state to steady state. We have also confirmed that the metacommunity exhibits either 5-cluster or 6-cluster for a random choice of any two patches. Due to predation turnoff, one of ten patches goes to a very low population density forming a source-sink dynamics (Fig. 2b and 2c). In this case, there exists only one sink patch. In addition, depending on the network configuration (i.e., ring network with different choices of predation turnoff), two sink patches can also exist in the 5-cluster and 6-cluster states.

Using a similar approach, we extend the predation turnoff in three patches and explore the corresponding dynamics. Since one can choose three random patches in many possible ways, we present three specific cases in Fig. 3 with a schematic and its corresponding time series. If three chosen patches are successive in the ring network (i.e., *i, i* + 1 and *i* + 2), then the metacommunity dynamics for predation turnoff in those patches lead to a 6-cluster. Note that we use patch-wise identical initial conditions (i.e., 1-cluster). In similar to previous sections, 1-cluster is changed into a 6-cluster, but in terms of steady states. The steady states in Figs. 2(c) and 3(a) are at different population densities despite they exhibit 6-cluster. If three patches are chosen in such a way that the second patch is alternative to the other chosen patches (i.e., *i, i* − 2, and *i* + 2), then the metacommunity exhibits a 5-cluster (see Fig. 3(b)). Though the number of predation turnoff is different in Figs. 3(b)) and 2(b), metacommunity still exhibits 5-cluster. As a third case, we present a 9-cluster for predation turnoff in three patches (Fig. 3(c)) if two patches are successive and the third one is two patches away (i.e., patch index *i, i* + 1 and *i* + 4). Moreover, there exist other clusters such as 8-cluster and 10-cluster and also other network configurations for predation turnoff in three patches.

**Figure 3:**
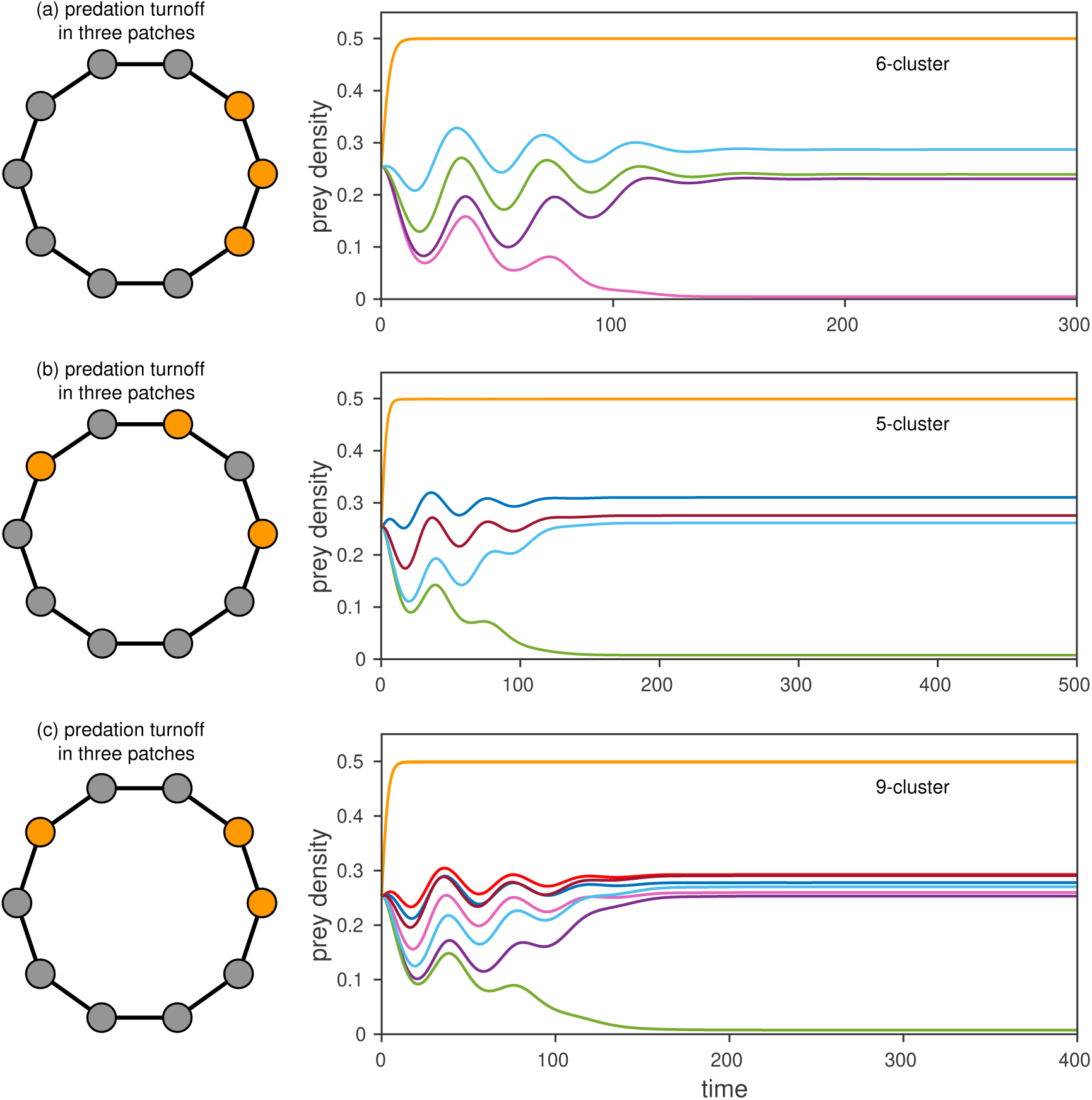
Metacommunity dynamics for predation turnoff in three patches: Considering a synchronized oscillatory state (1-cluster) in 10 patches, we show the schematic representation of predation turnoff and the corresponding dynamics in the left and right panel, respectively. Metacommunity exhibits (a) 6-cluster for predation turnoff in three consecutive patches, (b) 5-cluster for predation turnoff in three alternative patches, and (c) 9-cluster for predation turnoff in three patches in which two patches are chosen consecutively, and third patch is chosen two patch away from the consecutive patches. Here the parameter values are *r* = 0.5, *k* = 0.5, *b* = 0.16, *e* = 0.5, *m* = 0.2, *d*_*v*_ = 0.001, and *d*_*h*_ = 1.3.

Using a synchronized oscillatory (1-cluster) state as our initial assumption, we addressed the dynamics for predation turnoff in one, two and three patches in Figs. 2 and 3. This raises a question: can metacommunity exhibit the similar qualitative dynamics for an increasing number of predation turnoff and even for predation turnoff in all patches? To address this question, we summarize the dynamics in Fig. 4 as a function of predation turnoff with the increasing number of patches. Since there exist multiple cluster sizes for the predation turnoff in *j* number of patches, we collect all the possible cluster sizes using 100 simulations in which patches are chosen randomly. Figure 4 confirms both 5- and 6-clusters for predation turnoff in one and two patches as observed in Figs. 2. Similarly, for predation turnoff in three patches, we get 8- and 10-cluster along with 5-, 6-, and 9-clusters (as observed in Fig. 3). As an increase in the number of patches where the predation is turned off, the metacommunity exhibits other cluster sizes with unique possibilities. For example, predation turnoff in four patches leads to 3-cluster along with 5-, 6-, 9-, and 10-clusters. From the cluster sizes distributed horizontally in Fig. 4, one can identify that a specific cluster can exist for a different number of predation turnoff. For example, 4-cluster can exist when the predation turnoff from two to five patches. On the other hand, the cluster size distributed vertically determines the possible minimum and maximum cluster sizes for a specific number of predation turnoff. For example, the 3-cluster is the minimum one that the metacommunity can exhibit for predation turnoff in four patches, and this cluster is unique because it does not exist for any other choices of predation turnoff. Moreover, there exists a transition to 1-cluster for a higher number of predation turnoff. For example, predation turnoff in more than 5 patches leads to 1-cluster steady state indicating the critical number of patches subject to predation turnoff. Above this critical number, the predator in all patches goes to extinction. Below the critical value, metacommunity exhibits multicluster depending on the network configuration, and thus promoting the metacommunity persistence. The mechanism behind all these multi-clusters is that predation turnoff strongly alters the the connectivity strength between patches.

**Figure 4:**
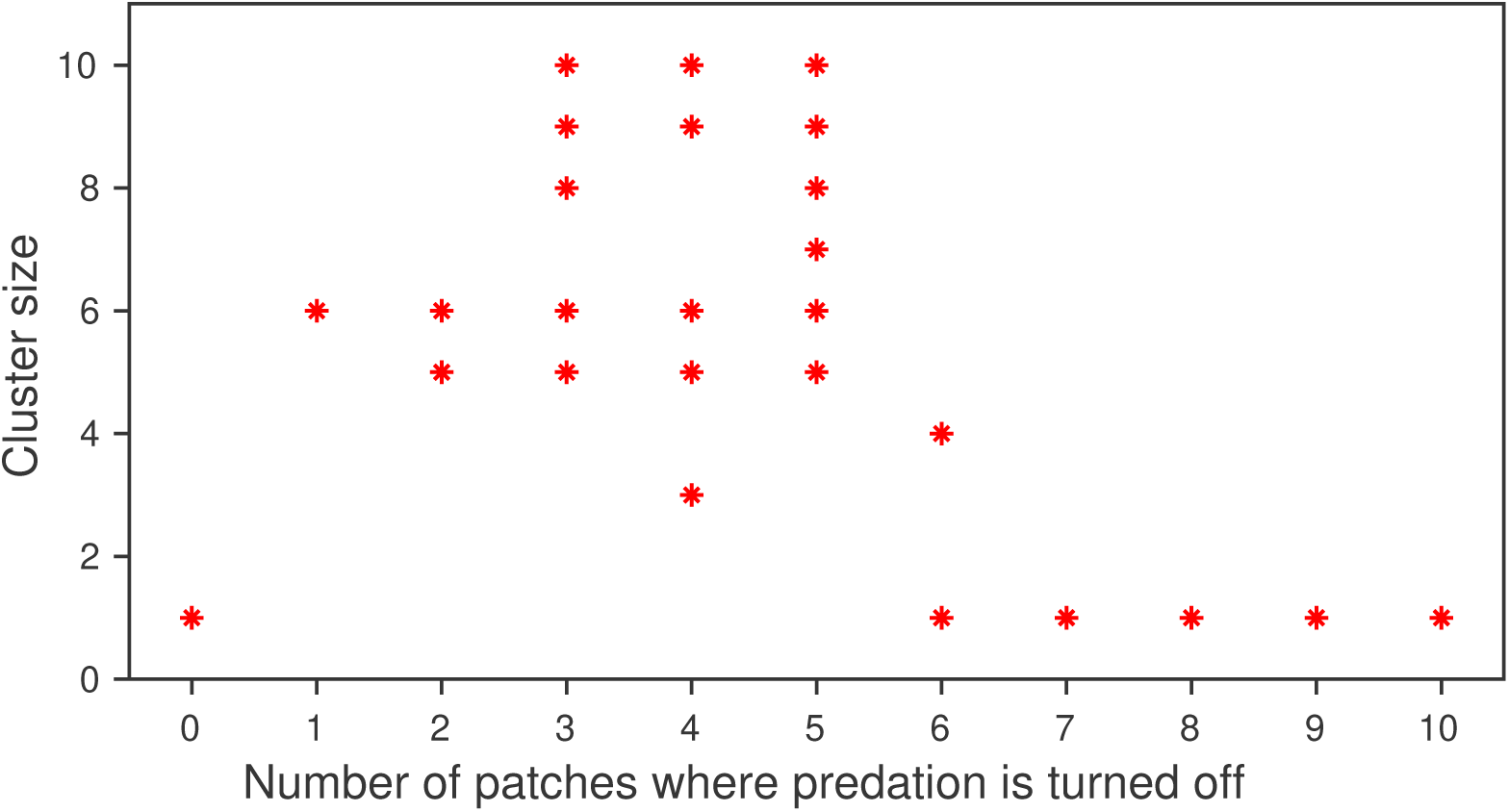
Metacommunity dynamics for predation turnoff in increasing number of patches: Cluster size is shown for increasing number of predation turnoff considering 1-cluster before applying predation turnoff. Predation turnoff in more than five patches leads to 1-cluster steady state with predators extinction. Here the parameter values are *r* = 0.5, *k* = 0.5, *b* = 0.16, *e* = 0.5, *m* = 0.2, *d*_*v*_ = 0.001, and *d*_*h*_ = 1.3.

### 3.2. Predation turnoff in source-sink dynamics

It is clear from the previous section that predation turnoff in selective patches changes the synchronized oscillatory state into asynchronized (multicluster) states. However, metacommunity without predation turnoff can even exhibit asynchronized (inhomogeneous) states depending on the initial conditions. In this section, considering an inhomogeneous (i.e., multicluster) state as a starting point of our analysis, we present how predation turnoff affects the metacommunity persistence. In Fig. 5(a) and 5(b), we show the metacommunity dynamics without and with predation turnoff, respectively. Here the schematic representation of ten patches is in the left panel, while their corresponding dynamics in the right panel. For the ring network topology, Figure 5(a) shows a 3-cluster with source-sink dynamics. From ten patches distributed in 3-cluster, four patches form a cluster with a very low density indicating the sink in the metacommunity, whereas the other six patches form two different clusters with higher density indicating the source dynamics. Indeed, dispersal creates inhomogeneous states with a source-sink behaviour. Moreover, one can identify from the time series in Fig. 5(a) that two patches have the highest density forming a cluster.

**Figure 5:**
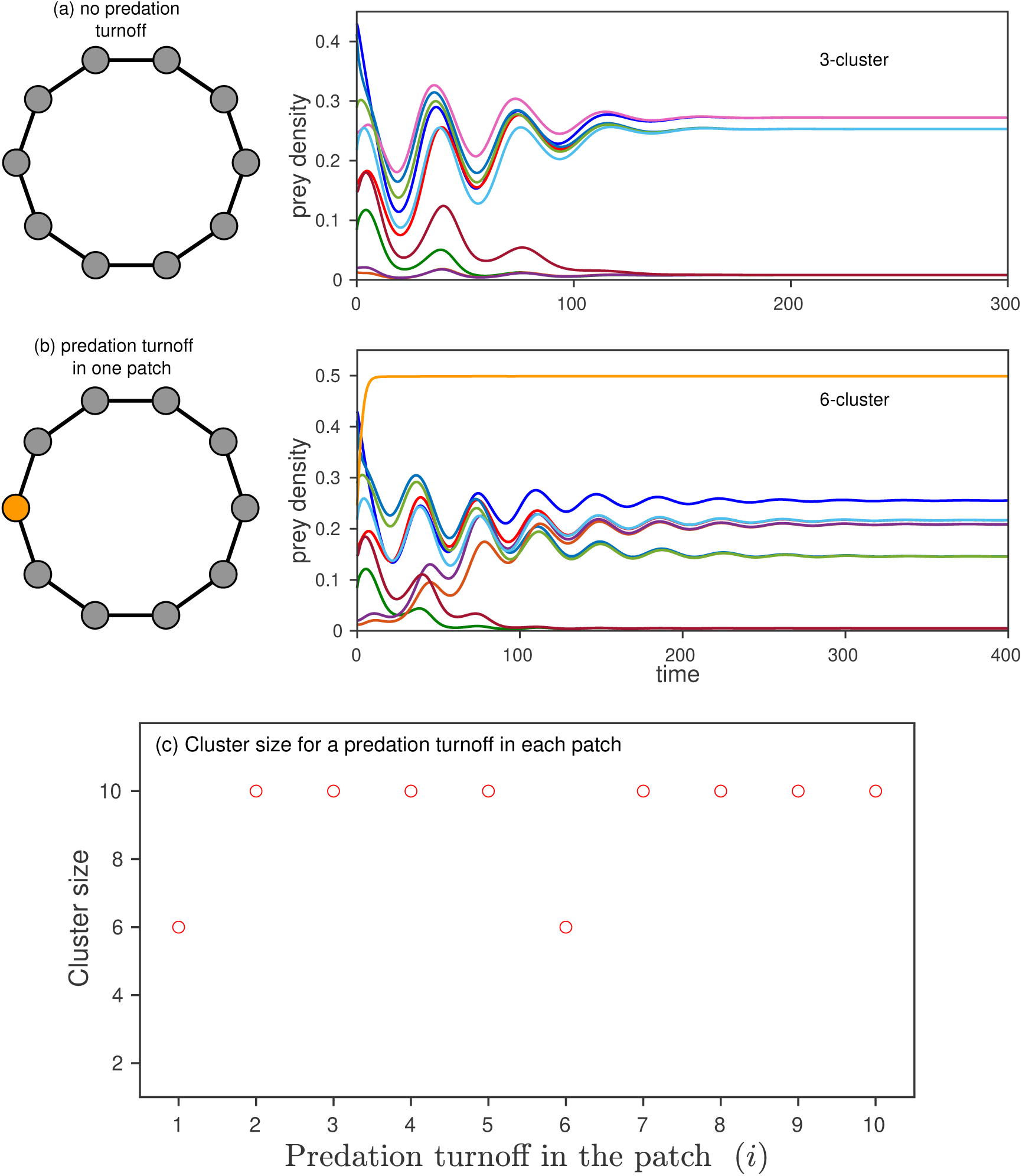
Metacommunity dynamics for predation turnoff in one patch considering an inhomogeneous state (3-cluster): Schematic representation of predation turnoff and the corresponding dynamics in the left and right panel, respectively. Metacommunity exhibits (a) 3-cluster when there is no predation turnoff, (b) 6-cluster for predation turnoff in a source patch. (c) Cluster size for a predation turnoff in the individual patch (i.e., patch-wise predation turnoff). Here the parameter values are *r* = 0.5, *k* = 0.5, *b* = 0.16, *e* = 0.5, *m* = 0.2, *d*_*v*_ = 0.001, and *d*_*h*_ = 1.3.

Now we apply the predation turnoff in a randomly chosen patch and compute the resulting cluster. In similar to previous sections, we observe a change in the clusters for predation turnoff (see Fig. 5(b)). In particular, 3-cluster is changed into a 6-cluster. Also, the prey density from the patch where the predation is turned off goes to a saturating state near the carrying capacity. One can choose a patch randomly, but we consider a source patch where the prey density is really higher (see Fig. 5(a)). Indeed, predation turnoff in this source patch reduces the number of sink. For example, four sink patches exist in Fig. 5(a) whereas only two sinks exists in Fig. 5(b). Further, it alters the density of other source patches as well as sink patches. On the other hand, one can find that predation turnoff in the sink changes only the number of clusters but does not reduce the number of sinks. Hence, depending on prey density in the selected patch, predation turnoff can lead to different clusters. Moreover, the 6-cluster that we observe in Fig. 5(b) is qualitatively different from the 6-cluster in Fig. 2(a) despite the predation turnoff is in only one patch in both cases. Indeed, the homogeneous and inhomogeneous population density of the metacommunity clearly distinguishes 6-cluster oscillatory and 6-cluster steady states for a predation turnoff in one patch.

Since the existing number of clusters corresponding to a predation turnoff depends on the quality of patches (i.e., source or sink), we compute the cluster size for patch-wise predation turnoff. Figure 5(c) shows the cluster size for predation turnoff in each of ten patches. In this case, only patch 1 and patch 6 exhibit 6-cluster, whereas other patches exhibit 10-cluster. Indeed, these patch 1 and patch 6 together earlier formed a cluster with the highest population in Fig. 5(a), which in turn leads to a 6-cluster for a predation turnoff as observed in Fig. 5(b). Hence predation turnoff in the source patch has a strong impact on the metacommunity dynamics by reducing the number of sink. On contrary, predation turnoff in the sink patch leads to only higher number of cluster but does not reduce the number of sinks. Moreover, similar behaviour can be observed if we choose other initial conditions that exhibit a 3-cluster. On the other hand, even if we assume a 6-cluster inhomogeneous state as our initial dynamics in the ring network topology, we get more complex dynamical responses in the metacommunity.

We have shown the effect of predation turnoff in Figures 2-5 using 1-cluster and multi-cluster. To show the robustness of our results for predation turnoff in randomly chosen patches and even for a higher number of predation turnoff, we do the following: At first, we collect 1000 initial conditions that exhibit only 3-cluster when there is no predation turnoff. For each initial condition, we choose three patches randomly and then apply the predation turnoff. Subsequently, we collect the resulting network configuration, the corresponding cluster size and represent them in a bar chart. Figure 6 shows the percentage of randomly chosen three patches that exhibit a specific cluster over 1000 simulations. It is clear from Fig. 6 that (i) a higher size always exists for predation turnoff in three patches though we start with a 3-cluster state before predation turnoff, (ii) 10-cluster exists with higher probability, and (iii) each network configuration describing predation turnoff in selective patches exhibits a unique cluster. Though there exist more than one network configuration for a specific cluster, we show one network configuration on the top of the bar chart. For example, if the predation turnoff is in three consecutive patches, then 5-cluster is the only possible dynamics. Since we use always a ring-network topology, each network configuration of predation turnoff is independent of patch indices (i.e., three consecutive patches such as 9, 10 and 1 would exhibit the 5-cluster and the patches 3, 4 and 5 would exhibit the same 5-cluster). Also, if the patches for predation turnoff are chosen with one alternative to the other, then there exists a 5-cluster. Thus, the predation turnoff in randomly selected patches also promotes metacommunity persistence. The dynamics that we showed here still exist in the other network topologies (see the supplementary material for a fully-connected network and a random network).

**Figure 6:**
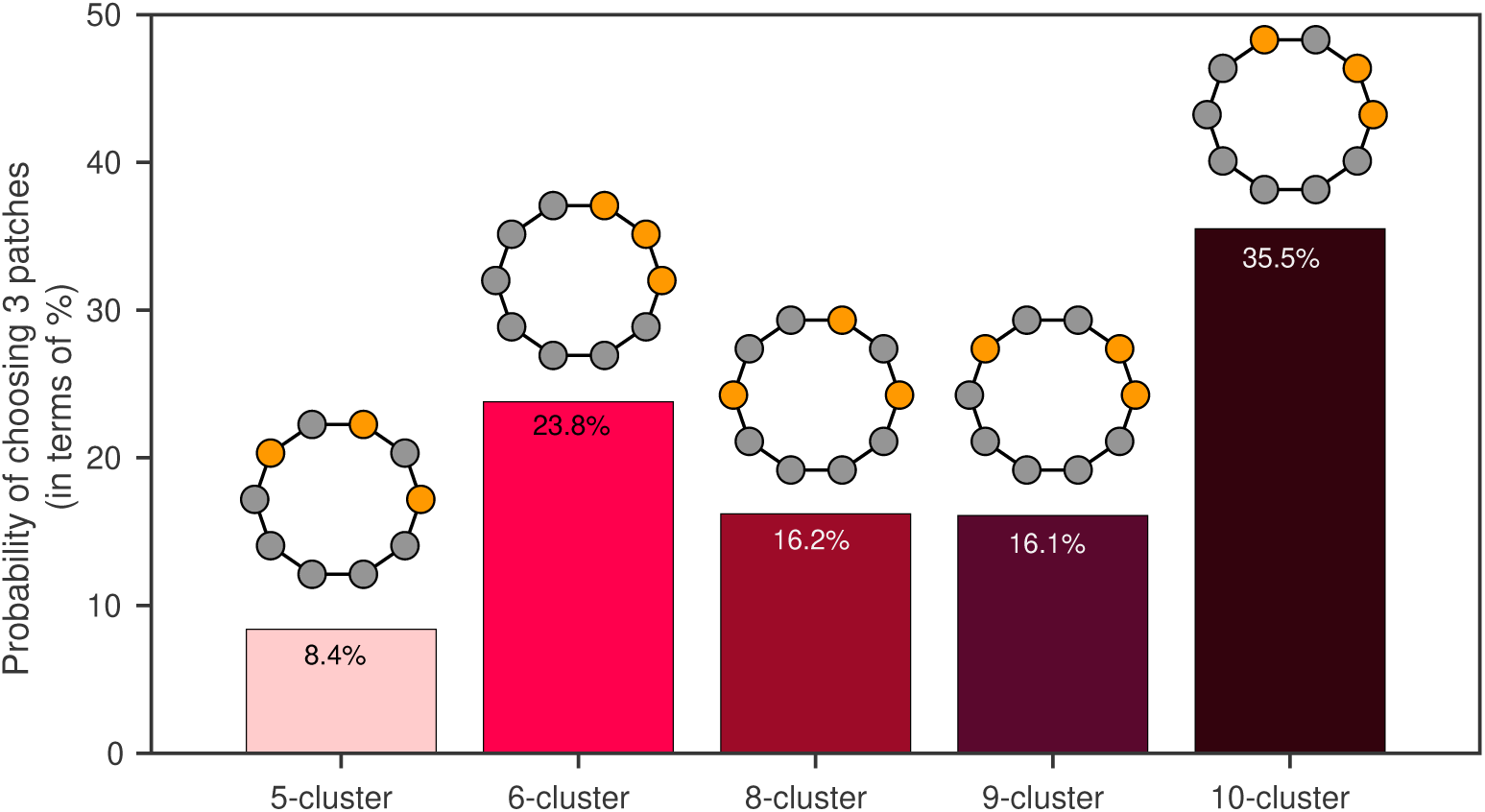
Metacommunity dynamics for predation turnoff in three random patches: We use 1000 initial conditions that exhibit 3-cluster in the ring network topology. For each initial condition, we apply predation turnoff in three randomly chosen patches and compute the cluster size. Each bar denotes the probability of obtaining a specific cluster from 1000 initial conditions. For each cluster, one network configuration of predation turnoff is shown on top of the bar chart. Here the parameter values are *r* = 0.5, *k* = 0.5, *b* = 0.16, *e* = 0.5, *m* = 0.2, *d*_*v*_ = 0.001, and *d*_*h*_ = 1.3.

### 3.3. Transition between clusters for predation rate variation

In the previous sections, we have presented the metacommunity dynamics for an abrupt change in the predation rate. Instead of an abrupt change, here we analyse how a continuous change in the predation itself can affect the metacommunity persistence. For example, the predation rate (*c*_*i*_) is varied identically in the patches where the predation was turned off earlier. This variation of predation includes the predation turnoff also as a specific case, for example, *c*_*i*_ = 0, for some *i*-th patch. Four different cases are taken into account, we present the dynamics for the predation variation in Fig. 7. Specifically, in three cases, we use the nearest neighbor dispersal (ring network) with predation variation in one, two and all patches. In the fourth case, we use a fully-connected (mean-field) network with predation variation in all patches. In each case, we start with a specific cluster with a predation rate *c*_*i*_ = 1 (filled circle in Fig. 7), and analyze the effect of predation variation in the considered range [0 1] (open circles in Fig. 7). Note that other patches have a fixed predation rate.

**Figure 7:**
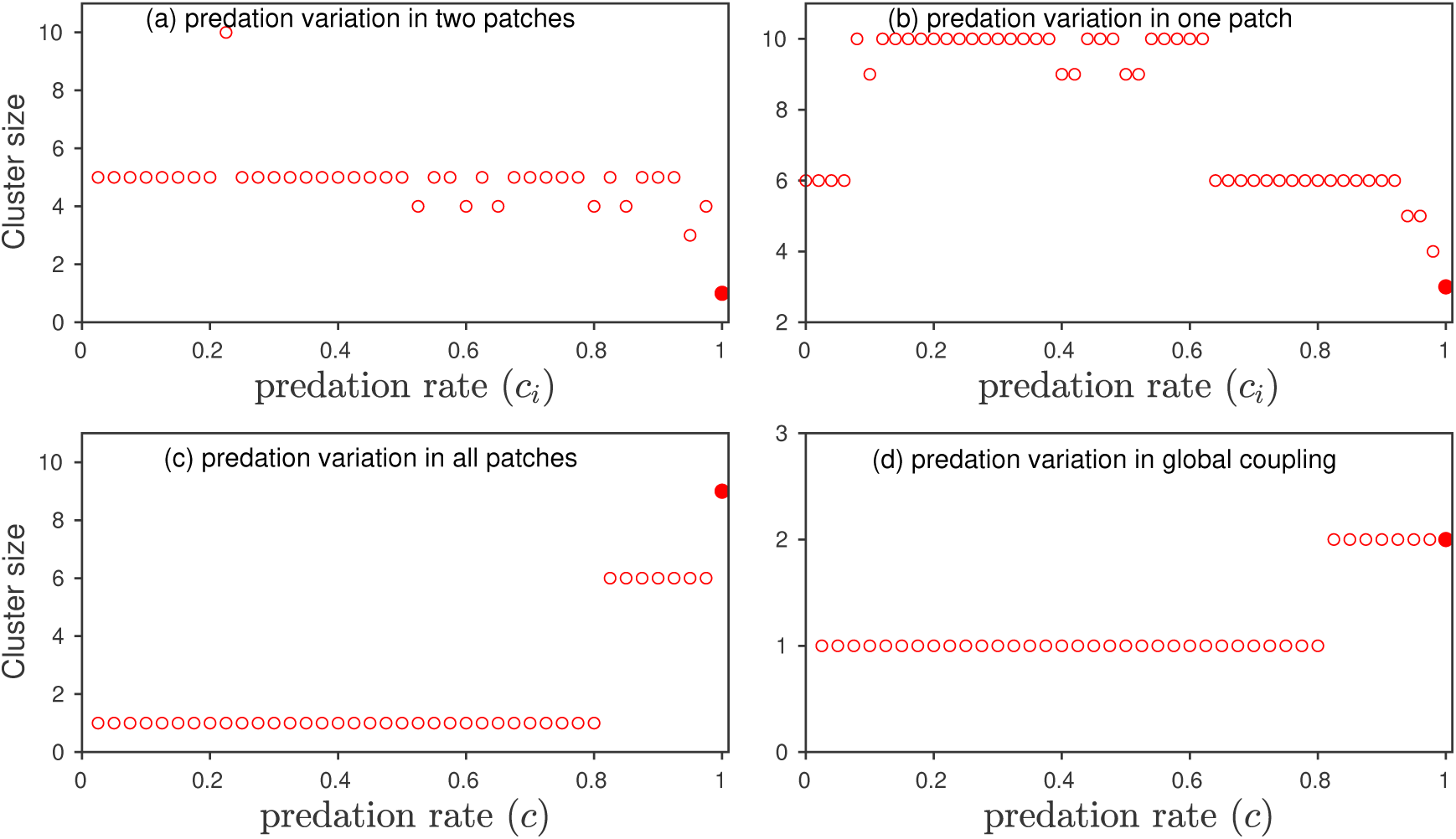
Cluster size as a function of predation rate: Predation rate (*c*_*i*_) is varied (a) identically in two patches, (b) in only one patch, (c)-(d) identically in all the patches. We use the ring-network topology in (a)-(c), wheres in (d), we use a fully-connected network. Filled circle in each panel denotes the initial cluster size before decreasing the predation rate. Here the parameter values are *r* = 0.5, *k* = 0.5, *b* = 0.16, *e* = 0.5, *m* = 0.2, *d*_*v*_ = 0.001, and *d*_*h*_ = 1.3.

Considering an homogeneous state (i.e., 1-cluster) for *c*_*i*_ = 1, we present the dynamics for identically varying predation rate in two selected patches (see Fig. 7a). Identical predation variation in two patches leads to a change in the number of clusters. For example, metacommunity exhibits mostly 5- and 6-clusters for higher values of predation rates, whereas only 5-cluster exists for lower predation rates. Specifically, at *c*_*i*_ = 0 (representing the predation turnoff in two patches), the metacommunity exhibits 5-cluster (which is similar to Fig.2(b)). In the second case, initially 3-cluster is taken into account (filled circle in Fig. 7b), we present the dynamics for varying predation rate in only one patch. In this case, for very low values and for high values of predation rate, metacommunity exhibits 6-cluster, whereas 9- and 10-clusters exist for intermediate values of predation rate. In addition, there exists 3- and 4-clusters for some higher predation rates. Despite that the predation turnoff determines a specific cluster in Figs. 2-6, but the predation variation in the range, [0 1], manifests different clusters for a fixed initial condition.

Instead of varying predation rate in some selected patches, we vary the predation rate identically in all patches and present the corresponding dynamics in Fig. 7(c) and 7(d) for local coupling and global coupling, respectively. Starting with a 9-cluster in the ring network, identical predation variation in all patches leads to a change in the number of clusters as well as a transition between clusters (see in Fig. 7(c)). For example, metacommunity exhibits 3-,6- and 9-clusters for the predation rates in the range [0 1]. Cluster size decreases for a decrease in the predation rate. There exists a critical predation rate for a transition from one cluster to another cluster. For example, 9-cluster becomes 6-cluster at *c* = 0.975, and 6-cluster is switched to 1-cluster at *c* = 0.8. In the case of fully-connected network (for the coupling equation, see the supplementary material), we start with an inhomogeneous state (i.e., 2-cluster) and present dynamics at different predation rates (see Fig. 7(d)). In similar to Fig. 7(c), there exists a transition between 2-cluster and 1-cluster at the predation rate *c* = 0.8125. Moreover, it is clear from Fig. 7 that cluster size fluctuates for varying predation rate in one or two patches, whereas a unique transition between clusters occurs for varying predation rate identically in all patches.

We extend our approach of predation rate variation by adding randomness to the predation rate of all patches. In this case, the predation rate of each *i*-th patch (*c*_*i*_) is chosen randomly, but they have a mean and a standard deviation (*σ*). We present the dynamics as a function of mean predation rate for four different standard deviation (*σ*) in Fig. 8 using a fixed initial condition. For the mean predation rate in absence of standard deviation (*σ* = 0), the dynamics is similar to Fig. 7(c) where transition between clusters take place at the critical value. By increasing the standard deviation, we observe multi-cluster for higher values of mean predation rates. Indeed, a small standard deviation changes the critical predation rate for the transition between 6-clusters and 1-cluster as observed in Fig. 7(c). Moreover, for all standard deviations, 1-cluster always exist for low values of mean predation rates. Indeed, there exists a critical predation rate below which 1-cluster exists and above which multicluster exists.

**Figure 8:**
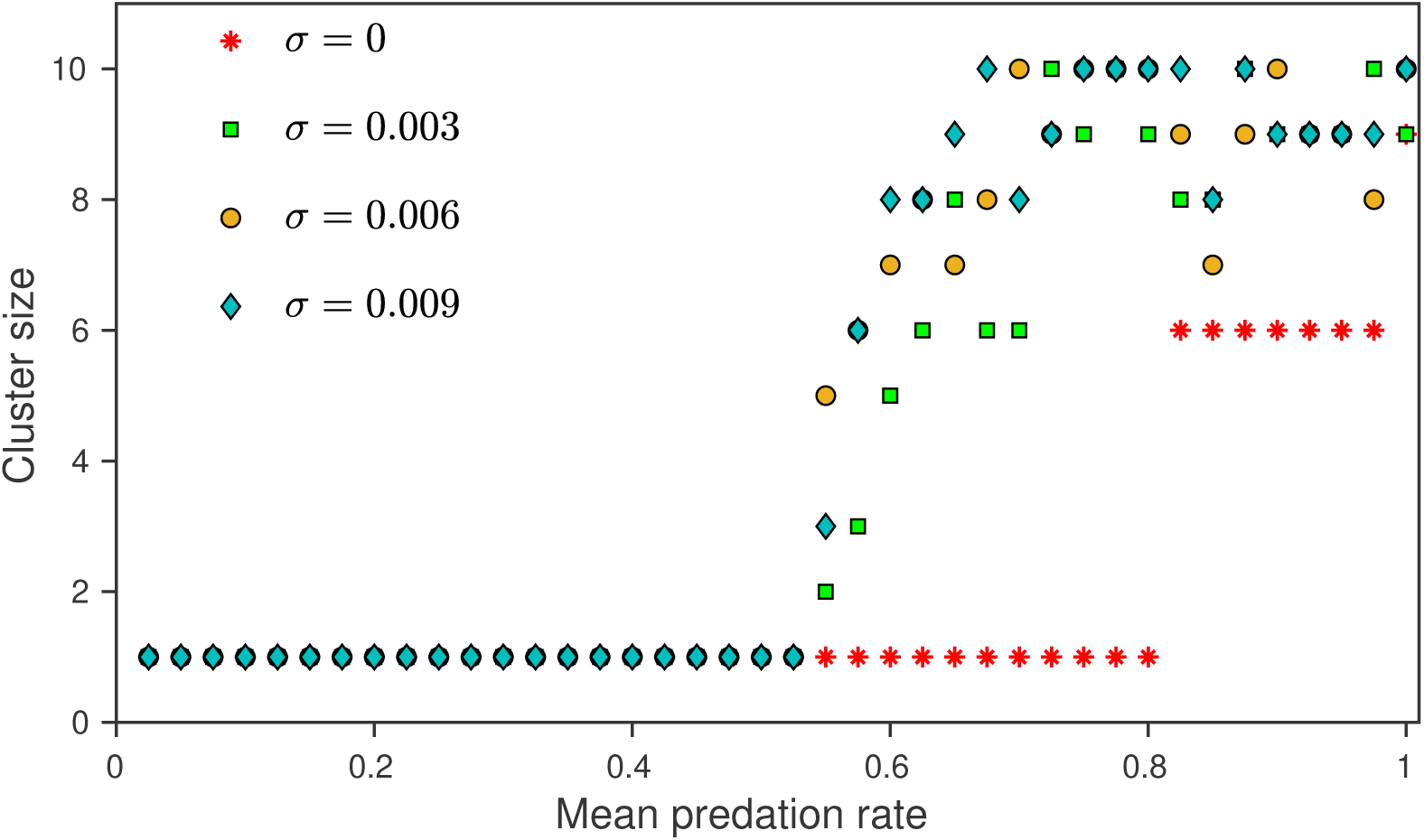
Cluster size as a function of mean predation rate: In the metacommunity, the predation rate of each patch (*c*_*i*_) is chosen randomly with a mean and a standard deviation (*σ*). There exists a critical mean predation rate that shows a transition between 1-cluster and multi-clusters for all standard deviations. Here the parameters are same as in Fig. 2.

### 3.4. Spatial-temporal dynamics for predation turnoff

To show the robustness of our results for a larger network structure, we use a network consisting of 50 patches and study the effect of predation turnoff on the metacommunity dynamics. Since the predator-prey metacommunity without predation turnoff can exhibit multiple dynamical states that include homogeneous and inhomogeneous states, we consider a specific inhomogeneous state (i.e., a 20-cluster) and study the effect of predation turnoff in three patches. We present the spatial-temporal dynamics of the metacommunity without and with predation turnoff in Fig. 9 with the top panels showing the spatial pattern, whereas the bottom panels show their corresponding time series. When there is no predation turnoff in the metacommunity, the spatial and temporal dynamics of 20-cluster state are shown in Fig. 9(a). In this 20-cluster state, prey densities in some patches are low indicating the source-sink dynamics. Now applying predation turnoff in three patches lead to the following changes in the dynamics: (i) a 20-cluster is changed into a 36-cluster (see Fig. 9(b) for the spatial dynamics and the corresponding time series), (ii) the prey density in those three patches goes to higher density near the carrying capacity, and (iii) there is a change in the source-sink dynamics. In similar to previous sections, a higher cluster always exists for a finite number of predation turnoff. From ecological perspective, a higher cluster size has less extinction risk as compare to low cluster size. Moreover, similar qualitative dynamics occur even if we consider homogeneous (1-cluster) or other inhomogeneous states before predation turnoff. Thus, predation turnoff in a very few patches can increase the metacommunity persistence through an increase in the number of clusters.

**Figure 9:**
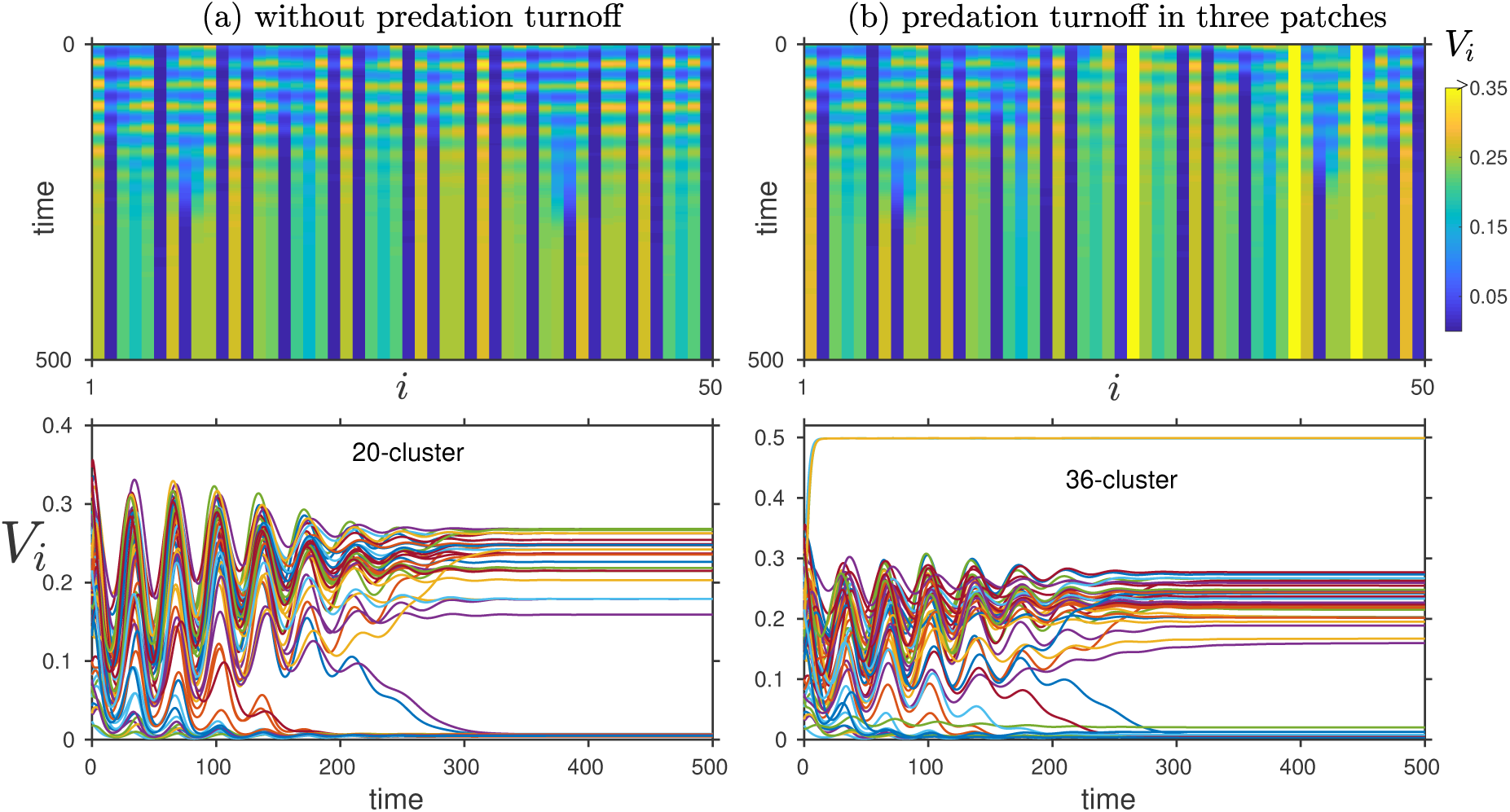
Metacommunity dynamics for a network of 50 patches: Left and right panels show the dynamics with and without predation turnoff, respectively. (a) Spatio-temporal dynamics with 20-cluster when there is no predation turnoff. (b) Spatio-temporal dynamics for predation turnoff in three patches exhibiting 36-cluster. Corresponding time series are shown in the bottom panels. Parameters are same as in Fig. 2.

## 4. Discussion

In this study, the interaction between local and spatial processes is taken into account, we have addressed how a predation turnoff in selective patches affect stability and persistence of a spatially distributed community. Using the clusters of a metacommunity, we have shown the impact of predation turnoff on synchronized, asynchronized and source-sink dynamics of the metacommunity. Our results reveal that predation turnoff can alter the synchronized dynamics into asynchronized dynamics by changing the 1-cluster oscillatory state into multi-cluster state. In a source-sink metacommunity, predation turnoff in a source patch reduces the number of sink patches and changes the number of clusters. In general, predation turnoff in finite number of patches can increase the number of clusters through inhomogeneous states, whereas predation turnoff in larger number can lead to complete extinction of predators. Indeed, our results determine the critical number of patches that can prevent local extinction through asynchronized (inhomogeneous) states. Subsequently, our findings emphasize the network configurations of predation turnoff that exhibit a unique cluster size. Moreover, prey density from the patches where predation is absent goes to a saturating state near the carrying capacity. In addition, our results in terms of patch-wise turnoff highlight the role of each patch on asynchronized (inhomogeneous) states. Thus, our findings through predation turnoff in selective patches reveal a stabilizing mechanism that can promote metacommunity persistence.

Predation is an important trophic interaction that shapes the community structure by changing species distributions. It is well known from numerous studies that predation can change the behavior, morphology, development, and abundance of prey (Werner and Peacor, 2003; Brown and Kotler, 2004). Though it reduces the prey abundance, it promotes diversity and also stabilizes the foodweb through top-down effect (Leibold, 1996; Carpenter and Kitchell, 1996; Pace et al., 1999). Theoretical studies on spatially distributed communities address that predation along with dispersal can promote persistence through asymmetric distributed population densities and source-sink dynamics (Jansen, 2001; Briggs and Hoopes, 2004). Moreover, recent experimental studies report that the presence of predators can alter fundamental mechanisms of community assembly and the scaling of diversity within metacommunities (Chase et al., 2009; Johnston et al., 2016). Since predation has significant impacts on various ecological processes, its removal or control can affect the stability of the system. Previous literature on predation removal shows the destabilizing effects through cascade of extinctions (Quince et al., 2005; Borrvall and Ebenman, 2006), but our study presents both positive and negative effects of predation turnoff in the spatially distributed metacommunity. In particular, our results highlight the positive effects through various multi-cluster patterns resulting from predation turnoff in only a finite number of patches, whereas cascading extinction of predators among patches occurs only when the predation turnoff is in more than half of patches of the network. Further, our results distinguish the dynamics between patchwise predation turnoff and for increasing number of predation turnoff. For example, predation turnoff in only one patch changes the perfectly synchronized oscillation into a phase-synchronized oscillation, whereas two patch predation turnoff changes the perfectly synchronized oscillation into multi-cluster steady states. Moreover, sinks are created in the metacommunity through predation turnoff. Further, our results highlight that synchronized and asynchronized dynamics strongly depend on both dispersal and the number of patches where the predation is turned off. Thus, our findings stresses the role of predation on metacommunity stability and persistence.

Population synchrony, as a general phenomenon of spatially distributed communities, has been ob-served at local and spatial scales (Hanski, 1998, 1999; Briggs and Hoopes, 2004; Liebhold et al., 2004). Dispersal, trophic interaction and environmental variation are three major reasons for the population synchrony (Bjørnstad et al., 1999; Koenig, 1999; Ims and Andreassen, 2000; Liebhold et al., 2004; Vasseur and Fox, 2009). Since synchrony might elevate a high risk of extinction, understanding the factors and mechanisms that affect the synchronized dynamics has received a greater attention in population ecology. As compare to synchronized behaviour, asynchrony among populations can increase the persistence by reducing the extinction risk. Dispersal plays a key role in the synchronized and asynchronized dynamics. By addressing the impact of predation on both synchronized and asynchronized dynamics, this study with a fixed dispersal reveals the strong implications of predation turnoff on the spatial metacommunity dynamics. For example, predation turnoff in only few patches can control the synchronized dynamics, which brings up a stabilizing mechanism to the metacommunity. Further, our results distinguish various dynamical behaviour depending on the number of predation turnoff which include a phase-synchronized oscillation, asynchronized steady states through various cluster size, and even homogeneous steady state. Though predation turnoff introduces a heterogeneity among the connected patches, but our results illustrate both homogeneous and heterogeneous behaviour depending on the number of patches where predation is absent. Thus, this study offers insights to predation-dispersal relationship.

One of the important dynamics of metacommunities resulting from dispersal is the source-sink behaviour described by the patch density with high and low abundance (Hanski, 1998; Amarasekare and Nisbet, 2001; Jansen, 2001; Gravel et al., 2010). Both synchronization and source-sink are the effects of dispersal, where the former leads to a high risk of extinction and the later can promote persistence through recolonization and rescue effect (Jansen, 2001; Gravel et al., 2010). Previous studies show that metacommunity without predation turnoff can still exhibit a source-sink behavior depending on dispersal and initial population densities. Indeed, our results are consistent with previous studies when there is no predation turnoff. Further, our study stresses the importance of predation turnoff on source-sink dynamics. For example, predation turnoff can change the synchronized population into a source-sink even for identical population densities (see Figs. 2-3). Subsequently, predation turnoff in the source-sink can change the number of sinks and increase the cluster size (see Figs. 5). In particular, predation turnoff in a source patch reduces the number of sinks, whereas predation turnoff in the sink does not alter the sink patches and only increases the cluster size. As habitat quality of the source can differ completely from that of the sink, our results explain the impact of predation turnoff in individual source and sink patches. Thus, our study highlights the role of predation on the source-sink dynamics.

In summary, this study presents that predation turnoff in some selective patches of the spatially distributed community has strong influence on the synchronized, asynchronized and source-sink dynamics. Using a simple predator-prey metacommunity model in a ring-network topology, we have addressed the impact of predation on metacommunity stability. But, more complex dynamics might occur if we use a metacommunity consisting of higher trophic interactions or even foodwebs, and also other network structures with different dispersal processes. Moreover, the dynamics that we observe might hold even if we include the environmental stochasticity in the metacommunity model, which requires a further investigation. Considering the predation removal as an evolutionary change or a climatic effect could have strong influence on the community stability and it requires a detailed investigation.

## Acknowledgments

DB acknowledges IISER-TVM for the financial support in terms of research fellowship. RA acknowledges IISER-TVM for the financial support to the postdoctoral research. VKC thanks DST, New Delhi for computational facilities under the DST-FIST programme (SR/FST/PS-1/2020/135) to the Department of Physics.

## References

Abbott KC. A dispersal-induced paradox: synchrony and stability in stochastic metapopulations. Ecology Letters 2011;14(11):1158–69. doi:10.1111/j.1461-0248.2011.01670.x.

Amarasekare P, Nisbet R. Spatial heterogeneity, source–sink dynamics, and the local coexistence of competing species. The American Naturalist 2001;158(6):572–84. doi:10.1086/323586.

Arumugam R, Dutta PS. Synchronization and entrainment of metapopulations: A trade-off among time-induced heterogeneity, dispersal, and seasonal force. Physical Review E 2018;97(6):062217. doi:10.1103/PhysRevE.97.062217.

Arumugam R, Dutta PS, Banerjee T. Dispersal-induced synchrony, temporal stability, and clustering in a mean-field coupled rosenzweig–macarthur model. Chaos 2015;25(10):103121. doi:10.1063/1.4933300.

Arumugam R, Sarkar S, Banerjee T, Sinha S, Dutta PS. Dynamic environment-induced multi-stability and critical transition in a metacommunity ecosystem. Phys Rev E 2019;99:032216. doi:10.1103/PhysRevE.99.032216.

Berger KM. Carnivore-livestock conflicts: Effects of subsidized predator control and economic correlates on the sheep industry. Conservation Biology 2006;20(3):751–61. doi:10.1111/j.1523-1739.2006.00336.x.

Bergstrom BJ, Arias LC, Davidson AD, Ferguson AW, Randa LA, Sheffield SR. License to kill: Reforming federal wildlife control to restore biodiversity and ecosystem function. Conservation Letters 2014;7(2):131–42. doi:10.1111/conl.12045.

Bjørnstad ON, Ims RA, Lambin X. Spatial population dynamics: analyzing patterns and processes of population synchrony. Trends in Ecology & Evolution 1999;14(11):427–32. doi:10.1016/S0169-5347(99)01677-8.

Borrvall C, Ebenman B. Early onset of secondary extinctions in ecological communities following the loss of top predators. Ecology letters 2006;9(4):435–42. doi:10.1111/j.1461-0248.2006.00893.x.

Brewer R. Principles of Ecology. Saunders (Philadelphia), 1979. Briggs CJ, Hoopes MF. Stabilizing effects in spatial parasitoid–host and predator–prey models: a review. Theoretical population biology 2004;65(3):299–315. doi:10.1016/j.tpb.2003.11.001.

Brown JS, Kotler BP. Hazardous duty pay and the foraging cost of predation. Ecology letters 2004;7(10):999–1014. doi:10.1111/j.1461-0248.2004.00661.x.

Caro T. Antipredator defenses in birds and mammals. University of Chicago Press, 2005. Carpenter SR, Kitchell JF. The trophic cascade in lakes. Cambridge University Press, 1996.

Chalcraft DR, Resetarits Jr WJ. Predator identity and ecological impacts: functional redundancy or functional diversity? Ecology 2003;84(9):2407–18. doi:10.1890/02-0550.

Chase JM, Biro EG, Ryberg WA, Smith KG. Predators temper the relative importance of stochastic processes in the assembly of prey metacommunities. Ecology letters 2009;12(11):1210–8. doi:10.1111/j.1461-0248.2009.01362.x.

Clark CW. Possible effects of schooling on the dynamics of exploited fish populations. ICES Journal of Marine Science 1974;36(1):7–14. doi:10.1093/icesjms/36.1.7.

Cott HB. Adaptive coloration in animals. London, Methuen & Co, 1940.

Crooks KR, Soulée ME. Mesopredator release and avifaunal extinctions in a fragmented system. Nature 2005;400:563–6. doi:10.1038/23028.

Dalla Rosa L, Secchi ER. Killer whale (orcinus orca) interactions with the tuna and swordfish longline fishery off southern and south-eastern brazil: a comparison with shark interactions. Journal of the Marine Biological Association of the United Kingdom 2007;87(1):135–40. doi:10.1017/S0025315407054306.

Dickman AJ. Complexities of conflict: the importance of considering social factors for effectively resolving human–wildlife conflict. Animal Conservation 2010;13(5):458–66. doi:10.1111/j.1469-1795.2010.00368.x.

Duffy JE. Biodiversity and ecosystem function: the consumer connection. Oikos 2002;99(2):201–19. doi:10.1034/j.1600-0706.2002.990201.x.

Earn DJ, Levin SA, Rohani P. Coherence and conservation. Science 2000;290(5495):1360–4. doi:10.1126/science.290.5495.1360.

Engen S, Sæther B. Generalizations of the moran effect explaining spatial synchrony in population fluctuations. The American Naturalist 2005;166(5):603–12. doi:10.1086/491690.

Goldwyn EE, Hastings A. When can dispersal synchronize populations? Theoretical population biology 2008;73(3):395–402. doi:10.1016/j.tpb.2007.11.012.

Goldwyn EE, Hastings A. The roles of the moran effect and dispersal in synchronizing oscillating populations. Journal of Theoretical Biology 2011;289:237–46. doi:10.1016/j.jtbi.2011.08.033.

Gore ML, Siemer WF, Shanahan JE, Schuefele D, Decker DJ. Effects on risk perception of media coverage of a black bear-related human fatality. Wildlife Society Bulletin 2005;33(2):507–16. doi:10.2193/0091-7648(2005)33[507:EORPOM]2.0.CO;2.

Gouhier TC, Guichard F, Gonzalez A. Synchrony and stability of food webs in metacommunities. The American Naturalist 2010;175(2):E16–34. doi:10.1086/649579.

Gravel D, Guichard F, Loreau M, Mouquet N. Source and sink dynamics in meta-ecosystems. Ecology 2010;91(7):2172–84. doi:10.1890/09-0843.1.

Gusset M, Swarner MJ, Mponwane L, Keletile K, McNutt JW. Human–wildlife conflict in northern botswana: livestock predation by endangered african wild dog lycaon pictus and other carnivores. Oryx 2009;43(1):67–72. doi:10.1017/S0030605308990475.

Hanski I. Metapopulation dynamics. Nature 1998;396(6706):41–9. doi:10.1038/23876.

Hanski I. Metapopulation Ecology. New York: Oxford University Press, 1999.

Hayward MW, Kerley GI. Fencing for conservation: Restriction of evolutionary potential or a riposte to threatening processes? Biological Conservation 2009;142(1):1–13. doi:10.1016/j.biocon.2008.09.022.

Heino M, Kaitala V, Ranta E, Lindström J. Synchronous dynamics and rates of extinction in spatially structured populations. Proceedings of the Royal Society: Biological Sciences 1997;264(1381):481–6. doi:10.1098/rspb.1997.0069.

Henschel P, Hunter LTB, Coad L, Abernethy KA, Mühlenberg M. Leopard prey choice in the congo basin rainforest suggests exploitative competition with human bushmeat hunters. Journal of zoology 2011;285(1):11–20. doi:10.1111/j.1469-7998.2011.00826.x.

Holland MD, Hastings A. Strong effect of dispersal network structure on ecological dynamics. Nature 2008;456(7223):792–4. doi:10.1038/nature07395.

Holyoak M, Leibold MA, Holt RD. Metacommunities: spatial dynamics and ecological communities. University of Chicago Press, Chicago, USA., 2005.

Howeth JG, Leibold MA. Predation inhibits the positive effect of dispersal on intraspecific and interspecific synchrony in pond metacommunities. Ecology 2013;94(10):2220–8. doi:10.1890/12-2066.1.

Ims RA, Andreassen HP. Spatial synchronization of vole population dynamics by predatory birds. Nature 2000;408(6809):194–6. doi:10.1038/35041562.

Ims RA, Steen H. Geographical synchrony in microtine population cycles: a theoretical evaluation of the role of nomadic avian predators. Oikos 1990;57:381–7. doi:10.2307/3565968.

Jackson RM, Wangchuk R. A community-based approach to mitigating livestock depredation by snow leopards. Human Dimensions of Wildlife 2004;9(4):1–16. doi:10.1080/10871200490505756.

Jansen VAA. The dynamics of two diffusively coupled predator–prey populations. Theoretical Population Biology 2001;59(2):119–31. doi:10.1006/tpbi.2000.1506.

Johnson CN, Wallach AD. The virtuous circle: predator-friendly farming and ecological restoration in australia. Restoration Ecology 2016;24(6):821–6. doi:10.1111/rec.12396.

Johnston NK, Pu Z, Jiang L. Predator identity influences metacommunity assembly. The Journal of animal ecology 2016;85(5):1161–70. doi:10.1111/1365-2656.12551.

Kendall BE, Bjørnstad ON, Bascompte J, Keitt TH, Fagan WF. Dispersal, environmental correlation, and spatial synchrony in population dynamics. The American Naturalist 2000;155(5):628–36. doi:10.1086/303350.

Koenig WD. Spatial autocorrelation of ecological phenomena. Trends in Ecology & Evolution 1999;14(1):22–6. doi:10.1016/S0169-5347(98)01533-X.

Korpimäki E, Norrdahl K. Experimental reduction of predators reverses the crash phase of small-rodent cycles. Ecology 1998;79(7):2448–55. doi:10.1890/0012-9658(1998)079[2448:EROPRT]2.0.CO;2.

Lande R, Engen S, Sæther BE. Spatial scale of population synchrony: environmental correlation versus dispersal and density regulation. The American Naturalist 1999;154(3):271–81. doi:10.1086/303240.

Leibold MA. A graphical model of keystone predators in food webs: trophic regulation of abundance, incidence, and diversity patterns in communities. The American Naturalist 1996;147(5):784–812. doi:10.1086/285879.

Lennox RJ, Gallagher AJ, Ritchie EG, Cooke SJ. Evaluating the efficacy of predator removal in a conflict-prone world. Biological Conservation 2018;224:277–89. doi:10.1016/j.biocon.2018.05.003.

Liebhold A, Koenig WD, Bjørnstad ON. Spatial synchrony in population dynamics. Annu Rev Ecol Evol Syst 2004;35:467–90. doi:10.1146/annurev.ecolsys.34.011802.132516.

Lima SL, Dill LM. Behavioral decisions made under the risk of predation: a review and prospectus. Canadian journal of zoology 1990;68(4):619–40. doi:10.1139/z90-092.

Löe J, Röskaft E. Large carnivores and human safety: a review. AMBIO: a journal of the human environment 2004;33(6):283–8. doi:10.1579/0044-7447-33.6.283.

Loreau M, Naeem S, Inchausti P. Biodiversity and Ecosystem Functioning: Synthesis and Perspectives. Oxford University Press, 2002.

McCann K, Hastings A, Huxel GR. Weak trophic interactions and the balance of nature. Nature 1998;395(6704):794–8. doi:10.1038/27427.

Menge BA, Olson AM. Role of scale and environmental factors in regulation of community structure. Trends in Ecology & Evolution 1990;5(2):52–7. doi:10.1016/0169-5347(90)90048-I.

Menge BA, Sutherland JP. Community regulation: variation in disturbance, competition, and predation in relation to environmental stress and recruitment. The American Naturalist 1987;130(5):730–57. doi:10.1086/284741.

Mishra C. Livestock depredation by large carnivores in the indian trans-himalaya: conflict perceptions and conservation prospects. Environmental conservation 1997;24(4):338–43. doi:10.1017/S0376892997000441.

Moran PA. The statistical analysis of the canadian lynx cycle. ii. synchronization and meteorology. Australian Journal of Zoology 1953;1(3):291–8. doi:10.1071/ZO9530291.

Ogada MO, Woodroffe R, Oguge NO, Frank LG. Limiting depredation by african carnivores: the role of livestock husbandry. Conservation biology 2003;17(6):1521–30. doi:10.1111/j.1523-1739.2003.00061.x.

Oli MK, Taylor IR, Rogers ME. Snow leopard panthera uncia predation of livestock: an assessment of local perceptions in the annapurna conservation area, nepal. Biological Conservation 1994;68(1):63–8. doi:10.1016/0006-3207(94)90547-9.

Pace ML, Cole JJ, Carpenter SR, Kitchell JF. Trophic cascades revealed in diverse ecosystems. Trends in ecology & evolution 1999;14(12):483–8. doi:10.1016/S0169-5347(99)01723-1.

Paine RT. Food web complexity and species diversity. The American Naturalist 1966;100(910):65–75. doi:10.1086/282400.

Pech RP, Sinclair A, Newsome A, Catling P. Limits to predator regulation of rabbits in australia: evidence from predator-removal experiments. Oecologia 1992;89:102–12. doi:10.1007/BF00319021.

Penteriani V, del Mar Delgado M, Pinchera F, Naves J, Fernández-Gil A, Kojola I, Härkönen S, Norberg H, Frank J, Fedriani JM, et al. Human behaviour can trigger large carnivore attacks in developed countries. Scientific reports 2016;6(1):1–8. doi:10.1038/srep20552.

Quince C, Higgs PG, McKane AJ. Deleting species from model food webs. Oikos 2005;110(2):283–96. doi:10.1111/j.0030-1299.2005.13493.x.

Ranta E, Kaitala V, Lindström J, Helle E. The moran effect and synchrony in population dynamics. Oikos 1997;:136–42doi:10.2307/3545809.

Ranta E, Kaitala V, Lindström J, Linden H. Synchrony in population dynamics. Proceedings of the Royal Society of London Series B: Biological Sciences 1995;262(1364):113–8. doi:10.1098/rspb.1995.0184.

Reynolds JC, Tapper SC. Control of mammalian predators in game management and conservation. Mammal Review 1996;26(2-3):127–55. doi:10.1111/j.1365-2907.1996.tb00150.x.

Ripa J. Analysing the moran effect and dispersal: their significance and interaction in synchronous population dynamics. Oikos 2000;89(1):175–87. doi:10.1034/j.1600-0706.2000.890119.x.

Ritchie EG, Johnson CN. Predator interactions, mesopredator release and biodiversity conservation. Ecology letters 2009;12(9):982–98. doi:10.1111/j.1461-0248.2009.01347.x.

Rosenzweig ML, MacArthur RH. Graphical representation and stability conditions of predator-prey interactions. The American Naturalist 1963;97(895):209–23. doi:10.1086/282272.

Ryberg WA, Smith KG, Chase JM. Predators alter the scaling of diversity in prey metacommunities. Oikos 2012;121(12):1995–2000. doi:10.1111/j.1600-0706.2012.19620.x.

Satake A, Bjørnstad ON, Kobro S. Masting and trophic cascades: interplay between rowan trees, apple fruit moth, and their parasitoid in southern norway. Oikos 2004;104(3):540–50. doi:10.1111/j.0030-1299.2004.12694.x.

Shurin JB, Allen EG. Effects of competition, predation, and dispersal on species richness at local and regional scales. The American Naturalist 2001;158(6):624–37. doi:10.1086/323589.

Vasseur DA, Fox JW. Phase-locking and environmental fluctuations generate synchrony in a predator– prey community. Nature 2009;460(7258):1007–10. doi:10.1038/nature08208.

Wallach AD, Johnson CN, Ritchie EG, O’Neill AJ. Predator control promotes invasive dominated ecological states. Ecology letters 2010;13(8):1008–18. doi:10.1111/j.1461-0248.2010.01492.x.

Weise MJ, Harvey JT. Impact of the california sea lion (zalophus californianus) on salmon fisheries in monterey bay, california. Fishery Bulletin 2005;103(4):685–96.

Werner EE, Peacor SD. A review of trait-mediated indirect interactions in ecological communities. Ecology 2003;84(5):1083–100. doi:10.1890/0012-9658(2003)084[1083:AROTII]2.0.CO;2.

